# Roles of the 5-HT2C receptor on zebrafish sociality

**DOI:** 10.1101/2022.09.12.507567

**Authors:** Layana Aquino de Moura, Maryana Pereira Pyterson, Ana Flávia Nogueira Pimentel, Fernanda Araújo, Loanne Valéria Xavier Bruce de Souza, Caio Henrique Moura Mendes, Bruna Patrícia Dutra Costa, Diógenes Henrique de Siqueira-Silva, Monica Lima-Maximino, Caio Maximino

## Abstract

Serotonin (5-HT) receptors have been implicated in social behavior in vertebrates. Zebrafish (*Danio rerio*) have been increasingly being used behavioral neuroscience to study the neurobiological correlates of behavior, including sociality. Nonetheless, the role of 5-HT_2C_ receptors in different social functions were not yet studied in this species. Zebrafish were treated with the agonist MK-212 (2 mg/kg) or the antagonist RS-102221 (2 mg/kg) and tested in the social interaction and social novelty tests, conditional approach test, or mirror-induced aggressive displays. MK-212 increased preference for an unknown conspecific in the social investigation test, but also increased preference for the known conspecific in the social novelty test; RS-102221, on the other hand, decreased preference in the social investigation test but increased preference for the novel conspecific in the social novelty test. MK-212 also decreased predator inspection in the conditional approach test. While RS-102221 decreased time in the display zone in the mirror-induced aggressive display test, it increased display duration. Overall, these results demonstrate the complex role of 5-HT_2C_ receptors in different social contexts in zebrafish, revealing a participation in social plasticity in vertebrates.

## 1. Introduction

Zebrafish (*Danio rerio* Hamilton 1822) are capable of complex social behavior, including not only shoaling (Miller and Gerlai, 2012) but also the formation of social hierarchies (Paull et al., 2010), cooperation (Pimentel et al., 2021), and social learning (Lindeyer and Reader, 2010). Across vertebrates, social behavior is mediated by brain networks that include the lateral septum (ventral subpallium in zebrafish), extended medial amygdala (supracommissural and postcommisural subpallium in zebrafish), preoptic area and paraventricular nucleus, anterior and ventromedial hypothalami (ventral and anterior tuberal nuclei in zebrafish), and periaqueductal gray area (O’Connell and Hofmann, 2012). This network is involved in multiple forms of social behavior, including sexual behavior and courtship, aggression, and parental care, and its nodes are reciprocally connected (Goodson, 2005; O’Connell and Hofmann, 2011). These regions also receive extensive serotonergic projections (Soares et al., 2018b, 2018a), suggesting that this monoamine modulates different forms of social behavior.

In zebrafish, serotonin (5-HT) has been implicated mainly in defensive behavior (Lima-Maximino et al., 2020; Silva et al., 2018) and aggression (Barbosa et al., 2019; Filby et al., 2010; Theodoridi et al., 2017); however, there are some indications that 5-HT participates in social interactions in this species. In a social interaction test, in which preference was established between visual access to a single conspecific or an empty tank, treatment with buspirone, a 5-HT_1A_ receptor partial agonist, increases preference for a conspecific; in the social novelty test (in which preference is established for the original stimulus vs. a novel conspecific), however, no effect was observed (Barba-Escobedo and Gould, 2012); suggesting that this receptor is important for social preference, but not for processing social novelty. Treatment with acute fluoxetine, which blocks serotonin transporter and therefore increases 5-HT levels, decreases social preference for a shoal of conspecifics (Giacomini et al., 2016). Treatment with lysergic acid diethylamide (LSD), a non-selective agonist at 5-HT_2_ receptors (Appel et al., 2004), and 3,4-methylenedioxymethamphetamine (MDMA), which leads to serotonin transporter-mediated efflux of 5-HT (Hilber et al., 2005), decreases shoal cohesion in zebrafish (Green et al., 2012). In a version of the social preference test, in which the preference of a focal animal for the image of a shoal *vs*. the image of a single zebrafish with the *nacre* phenotype, MDMA increased social preference (Ponzoni et al., 2016); MDMA derivatives 2,5-dimetoxy-4-bromo-amphetamine hydrobromide (DOB) and *para*-methoxyamphetamine (PMA) showed the same effect, and the 5-HT_2A/2C_ antagonist ritanserin blocked the effects of these amphetamines (Ponzoni et al., 2016), suggesting a role for receptors of the 5-HT_2_ family in social preference. However, the specific involved receptor is not known. In mice expressing only the fully edited VGV isoform of the 5-HT_2C_ receptor, treatment with mCPP, a preferential 5-HT_2C_ agonist, decreases social interaction – an effect which is absent in wild-type mice, which nonetheless show decreased social interaction in relation to VGV animals (Martin et al., 2013). Moreover, most of the interactions displayed by VGV mice were aggressive, while wild-type animals displayed little aggression against conspecifics in that situation – suggesting that, at least in this case, RNA editing of the 5-HT_2C_ receptor is related to aggression (Martin et al., 2013).

The seemingly contradictory effects of MDMA in zebrafish – decreasing shoaling and increasing social preference – could also be due to differences in the underlying motivations of behavior in each test. Shoaling, for example, is usually interpreted as involving an antipredator component, as aversive stimuli increase shoal cohesion (Green et al., 2012; Miller and Gerlai, 2007). Nonetheless, shoaling also introduces potential costs, such as competition for resources, risks of injury from intraspecific conflict, and increased risk of disease transmission (Krause et al., 2007). Likewise, while social preference is usually interpreted mainly as driven by an appetitive/reinforcing drive, social interactions can also be aversive (Soares et al., 2018a), especially when novel conspecifics are involved (social novelty).

Another context which involves social decision-making under motivational conflict is predator inspection, a behavior in which an individual fish leaves a shoal to approach a potential predator (Dugatkin et al., 2010; Pimentel et al., 2021). In this context, the individual must decide between staying in the relative safety of the shoal and leave it to inspect the predator, therefore, increasing its own risk of predation. The probability of inspecting a predator is dependent on whether or not a conspecific will “reciprocate” by also inspecting, in a strategy called “conditional approach” (Dugatkin, 1988; Pimentel et al., 2019, 2021). In this context, predator inspection is usually understood as “cooperative”, “prosocial”, or “altruistic” behavior (Dugatkin, 1997). In both guppies (*Poecilia reticulata*) and zebrafish, it has been shown that individuals inspect predators but also show signs of fear-like behavior (Pimentel et al., 2019, 2021). Interestingly, acute treatment of guppies with fluoxetine, therefore increasing 5-HT brain levels, increases predator inspection as simultaneously increasing freezing, while the non-selective 5-HT receptor antagonist metergoline produced the opposite effect (Pimentel et al., 2019).

Aggressive interactions also involve motivational conflict, but this time between the benefits of winning a contest (e.g., accessing or maintaining access to resources, signaling fitness, etc.) and the risks associated with it. Moreover, aggressive interactions are intrinsically competitive, instead of cooperative. 5-HT has been proposed to mediate social plasticity in aggressive behavior, with its transient release (phasic-like responses) before and during fighting promoting aggressive motivation and offensive acts and inhibiting the motivation to flee, while its tonic release being negatively associated with the overall level of aggressiveness (de Boer et al., 2015). There is consistent evidence of a participation of 5-HT in the control of aggressive interactions in teleost fish. In the matrinxã *Brycon amazonicus*, the introduction of an intruder in a territory controlled by one resident individual increases 5-HT levels in the hypothalamus (Wolkers et al., 2015). In the African cichlid *Astalotilapia burtoni* – a species which displays joint territory defense –, aggression levels of neighbors, but not residents, is correlated with the expression of 5-HT_2C_ receptors in the dorsomedial telencephalon (Dm)(Weitekamp et al., 2017), an homolog of the mammalian associative amygdala system (Maximino et al., 2013). In the Siamese fighting fish *Betta splendens* (Dzieweczynski and Hebert, 2012; Eisenreich and Szalda-Petree, 2015; Forsatkar et al., 2013; Lynn et al., 2007) and in zebrafish, acute fluoxetine decreases mirror-induced aggressive displays (but see Barbosa et al., 2019, for phenotype-specific effects in zebrafish). When zebrafish are allowed to interact directly with a conspecific, however, fluoxetine decreases offensive aggression in dominants and eliminates freezing in subordinates (Theodoridi et al., 2017). The 5-HT_1A_ receptor appears to be involved, as treatment with the agonist 8-OH-DPAT decreased mirror-induced aggression in fighting fish (Clotfelter et al., 2007), while treatment with the antagonist WAY 100,635, increases offensive aggression against conspecifics in zebrafish (Filby et al., 2010). It has been proposed that these effects are mainly due to presynaptic action (de Boer et al., 2015). Taken together with results from both social preference/novelty experiments and conditional approach, 5-HT appears to reduce competitive-aggressive behavior through the 5-HT_1A_ receptor and promote cooperative-approach behaviors through 5-HT_2_ receptors.

The present work tests how 5-HT_2C_ receptors are involved in social interactions in zebrafish. Specifically, we propose that activating the 5-HT_2C_ receptor will increase social preference and conditional approach, but have not effect on mirror-induced aggression. Conversely, blocking 5-HT_2C_ receptors will decrease social preference and conditional approach, again with no effects in aggressive displays.

## 2. Methods

### 2.1. Animals and housing

123 zebrafish (*D. rerio*) from the longfin phenotype were used in the experiments; details for sample size calculations can be found on each experimental section, below. Outbred populations were used due to their increased genetic variability, decreasing the effects of random genetic drift that could lead to the development of uniquely heritable traits (Parra et al., 2009; Speedie and Gerlai, 2008). The populations used in the present experiments are expected to better represent the natural populations in the wild. Animals were bought from a commercial vendor (Fernando Peixes, Belém/PA) and arrived in the laboratory with an approximate age of 3 months (standard length = ± 1.4 mm), being quarantined for two weeks, before the experiment begin, when animals had an approximate age of 4 months (standard length = 23.0 ± 3.2 mm).

They were kept in mixed-sex tanks during acclimation, with an approximate ratio of 1:1 males to females (confirmed by body morphology). The breeder was licensed for aquaculture under Ibama’s (Instituto Brasileiro do Meio Ambiente e dos Recursos Naturais Renováveis) Resolution 95/1993. Animals were group-housed in 40 L tanks, with a maximum density of 25 fish per tank, for the aforementioned quarantine before experiments begun. Tanks were filled with non-chlorinated water at room temperature (28 °C) and a pH of 7.0-8.0. Lighting was provided by fluorescent lamps in a cycle of 14 hours of light-10 hours of darkness, according to standards of care for zebrafish (Lawrence, 2007). Other water quality parameters were as follows: hardness 100-150 mg/L CaCO3; dissolved oxygen 7.5-8.0 mg/L; ammonia and nitrite < 0.001 ppm. Potential suffering of animals was minimized by controlling for the aforementioned environmental variables and scoring humane endpoints (clinical signs, behavioral changes, bacteriological status), following Brazilian legislation (Conselho Nacional de Controle de Experimentação Animal - CONCEA, 2017). Animals were used for only one experiment and in a single behavioral test, to reduce interference from apparatus exposure. Experiments were approved by UEPA’s IACUC under protocol 06/18.

### 2.2. Drug treatments

Animals were treated either with vehicle (10% DMSO), the 5-HT_2C_ receptor agonist MK-212 (6-chloro-2-(1-piperazinyl)pyrazine), CPP; CAS Number 64022-27-1; 2 mg/kg, a dose shown to block alarm reactions in zebrafish (do Carmo Silva et al., 2021)), or the 5-HT_2C_ receptor antagonist RS-102221 (8-[5-(2,4-Dimethoxy-5-(4-trifluoromethylphenylsulphonamido)phenyl-5-oxopentyl]-1,3,8-triazaspiro[4.5]decane-2,4-dione; CAS Number 185376-97-0; 2 mg/kg, a dose shown to block post-alarm substance anxiety-like behavior in zebrafish (do Carmo Silva et al., 2021)). Experimenters were blind to drug treatment by coding vials, and animals were randomly assigned to treatment groups through the use of random numbers (http://jerrydallal.com/random/assign.htm). Drugs were dissolved in 10% DMSO, used as vehicle, and injected i.p. after cold-anesthesia (Kinkel et al., 2010; Matthews and Varga, 2011) in a volume of 5 µL. 30 min after drug or vehicle injection, animals were individually tested in the SI/SN, conditional approach, or MIA tests.

### 2.3. Experiment 1: Role of 5-HT2C receptors in social interaction and social novelty in zebrafish

After drug treatment, animals were individually netted and transferred to the experimental apparatus, a tank (13 × 17 × 13 cm [length × width × height]) positioned alongside two tanks with the same dimensions as the observation tank (Figure 1A). In this experiment, 30 animals were used, based on sample size calculation of the effects of buspirone on social interaction and social novelty (Barba-Escobedo and Gould, 2012). One animal was removed from the RS-102221 group after showing signs of altered health. Following the protocol by Barba-Escobedo and Gould (2012), a sequential design was used, with animals first exposed to the social interaction test (SI) and immediately exposed to the social novelty (SN) test. For the SI test, animals were allowed to freely explore the central tank for 10 min. before recording begun; during this acclimation period, visual contact with the adjacent tanks was blocked by white plastic sheets. After this period, the sheets were removed, allowing visual contact with both adjacent tanks, which contained either one conspecific (drawn from a different tank than the one that originated the focal animal) or an empty tank. The position of the conspecific (“stranger 1”) in either the leftmost or the rightmost tank was randomly varied for each subject in a counterbalanced fashion. Animals were then allowed to freely explore the tank for another 10 min. Sessions were filmed from the front using a digital camera (Samsung ES68, Carl Zeiss lens). Video recordings of the session were analyzed using TheRealFishTracker (https://www.dgp.toronto.edu/~mccrae/projects/FishTracker/); the following variables were analyzed:

**Figure 1.**
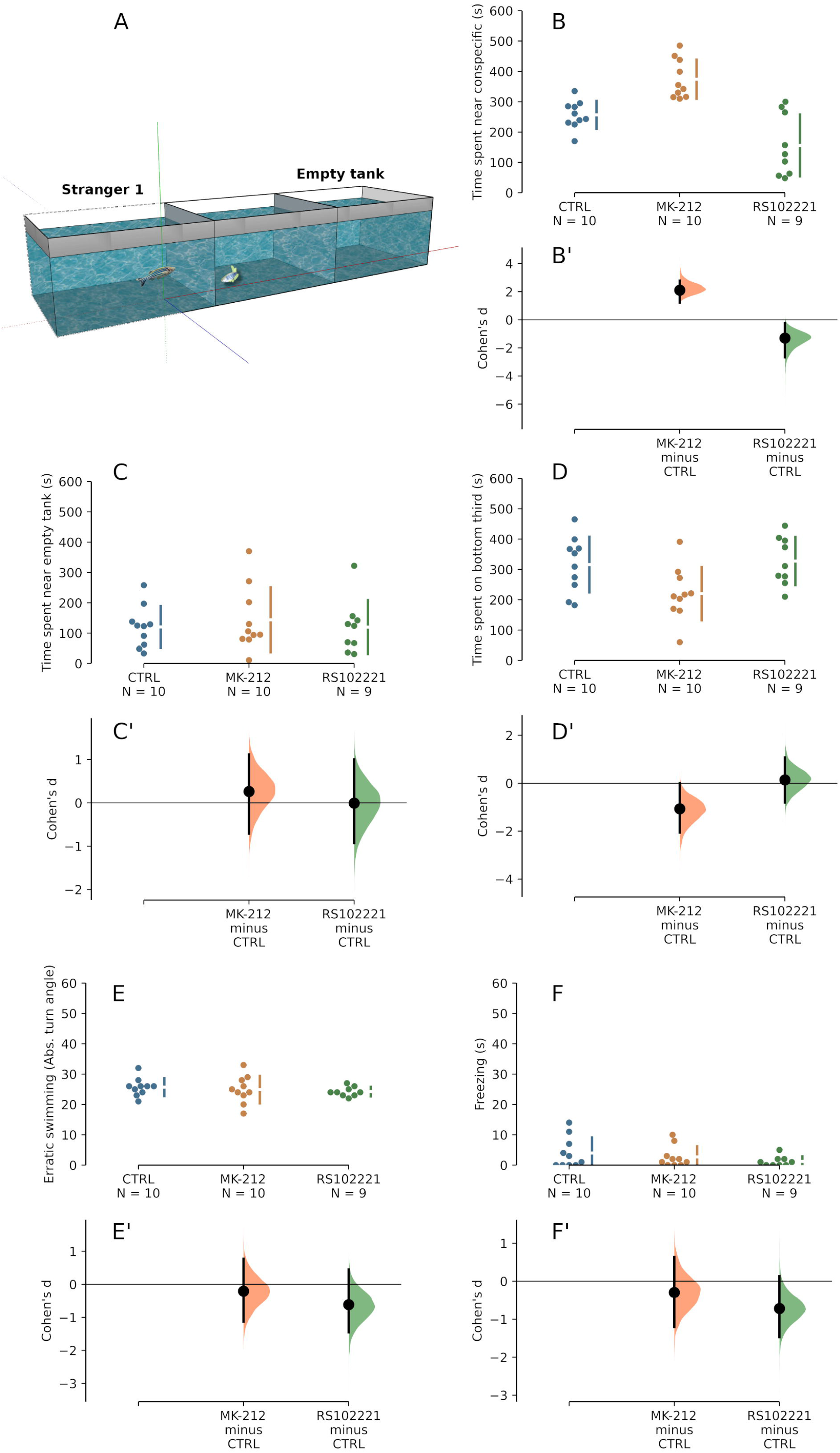
– In the social investigation test, MK-212 increases preference for a conspecific and RS-102221 decreases it. (A) Schematics of the test apparatus. (B) Raw data for the time spent in the section of the tank that is nearest to the conspecific. (B’) Mean differences between groups shown in (B), plotted as as bootstrap sampling distributions of Cohen’s *d* for treatments compared with controls. (C) Raw data for the time spent in the section of the tank that is nearest to the empty tank. (C’) Mean differences between groups shown in (C), plotted as as bootstrap sampling distributions of Cohen’s *d* for treatments compared with controls. (D) Raw data for the time spent in the bottom third of the tank. (D’) Mean differences between groups shown in (D), plotted as as bootstrap sampling distributions of Cohen’s *d* for treatments compared with controls. (E) Raw data for erratic swimming, measured as absolute turn angle. (E’) Mean differences between groups shown in (E), plotted as as bootstrap sampling distributions of Cohen’s *d* for treatments compared with controls. (F) Raw data for the time spent freezing (swimming speed < 0.5 cm/s). (F’) Mean differences between groups shown in (F), plotted as as bootstrap sampling distributions of Cohen’s *d* for treatments compared with controls. For all effect size plots, each mean difference (estimated Cohen’s *d*) is depicted as a dot. Each 95% confidence interval is indicated by the ends of the vertical error bars.

1. Time near stranger 1: The time spent in a X cm zone nearest to the conspecific;
2. Time far from stranger 1: The time spent in a X cm zone nearest to the empty tank;
3. Geotaxis: The time spent in the bottom third of the tank;
4. Erratic swimming: Absolute turn angle, in degrees.
5. Freezing: The time spent immobile (swimming speed < 0.5 cm/s)

Immediately after ending the SI trial, the plastic screens were lowered, and another conspecific (“stranger 2”, again drawn from a different tank than the one which originated the focal animal) was transferred to the empty tank (Figure 2A). Screens were raised, and the animal was allowed to freely explore the tank for another 10 min. The same variables were recorded for this stage.

**Figure 2.**
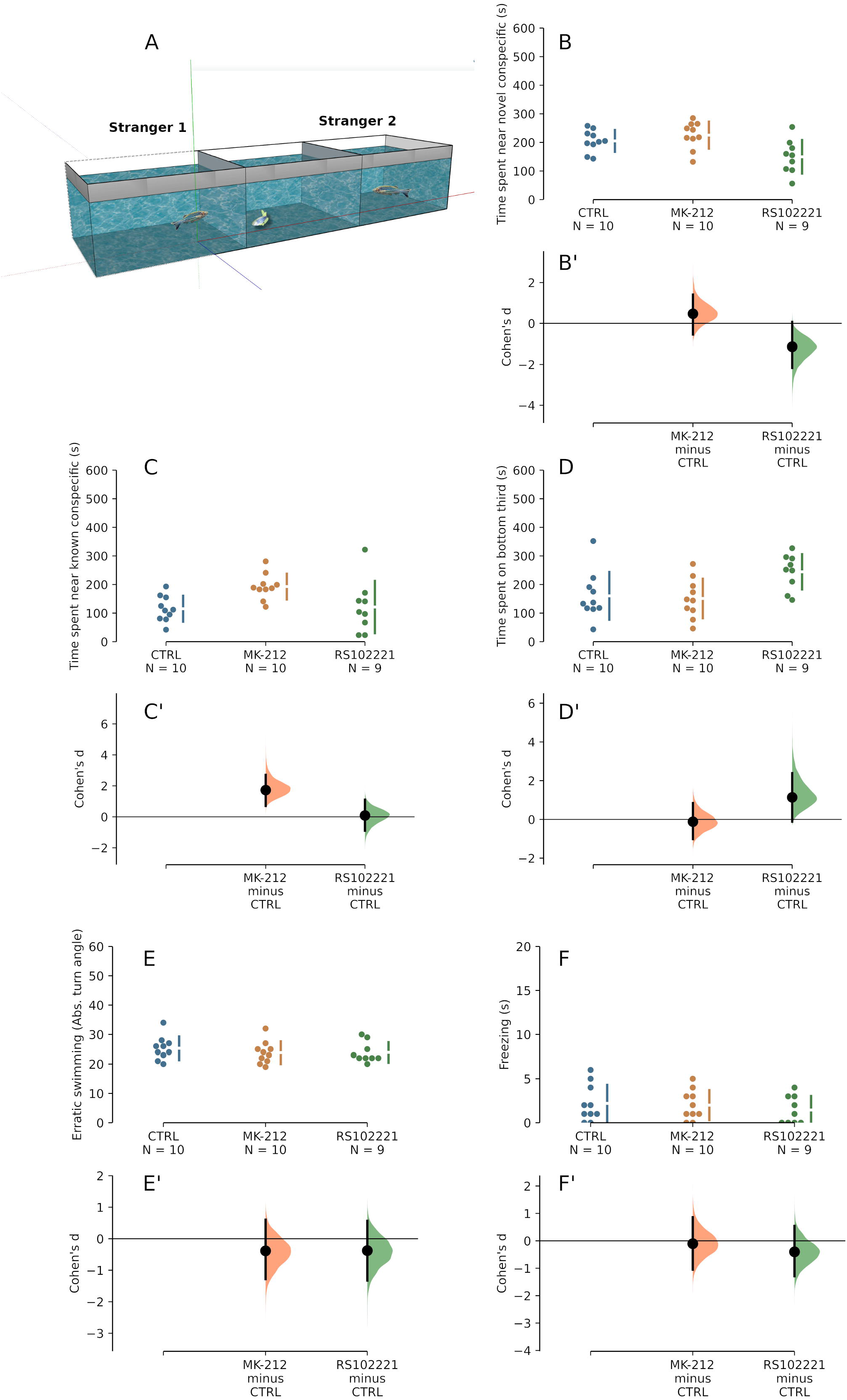
– In the social novelty test, MK-212 increases preference for a familiar conspecific (“stranger 1”) and RS-102221 decreases preference for a novel conspecific (“stranger 2”). (A) Schematics of the test apparatus. (B) Raw data for the time spent in the section of the tank that is nearest to the conspecific. (B’) Mean differences between groups shown in (B), plotted as as bootstrap sampling distributions of Cohen’s *d* for treatments compared with controls. (C) Raw data for the time spent in the section of the tank that is nearest to the empty tank. (C’) Mean differences between groups shown in (C), plotted as as bootstrap sampling distributions of Cohen’s *d* for treatments compared with controls. (D) Raw data for the time spent in the bottom third of the tank. (D’) Mean differences between groups shown in (D), plotted as as bootstrap sampling distributions of Cohen’s *d* for treatments compared with controls. (E) Raw data for erratic swimming, measured as absolute turn angle. (E’) Mean differences between groups shown in (E), plotted as as bootstrap sampling distributions of Cohen’s *d* for treatments compared with controls. (F) Raw data for the time spent freezing (swimming speed < 0.5 cm/s). (F’) Mean differences between groups shown in (F), plotted as as bootstrap sampling distributions of Cohen’s *d* for treatments compared with controls. For all effect size plots, each mean difference (estimated Cohen’s *d*) is depicted as a dot. Each 95% confidence interval is indicated by the ends of the vertical error bars.

### 2.4. Experiment 2: Role of 5-HT_2C_ receptors in conditional approach in zebrafish

The conditional approach strategy is observed when animals encounter a predator in the presence of conspecifics, eliciting predator inspection behavior. In this situation, if the conspecific reciprocates predator inspection, the probability of further inspection by both inspecting individuals is higher, suggesting cooperation (Dugatkin, 1997; Pimentel et al., 2021). In this experiment, 21 animals were used per group, based on a sample size calculation using the effect size of fluoxetine on conditional approach in guppies from Pimentel et al. (2019). After drug treatment, animals were transferred to the test tank, a 40.6 cm x 18 cm x 25 cm (length X width X height) glass tank whose bottom was divided in 10 equally-sized quadrants (Figure 3A).

**Figure 3.**
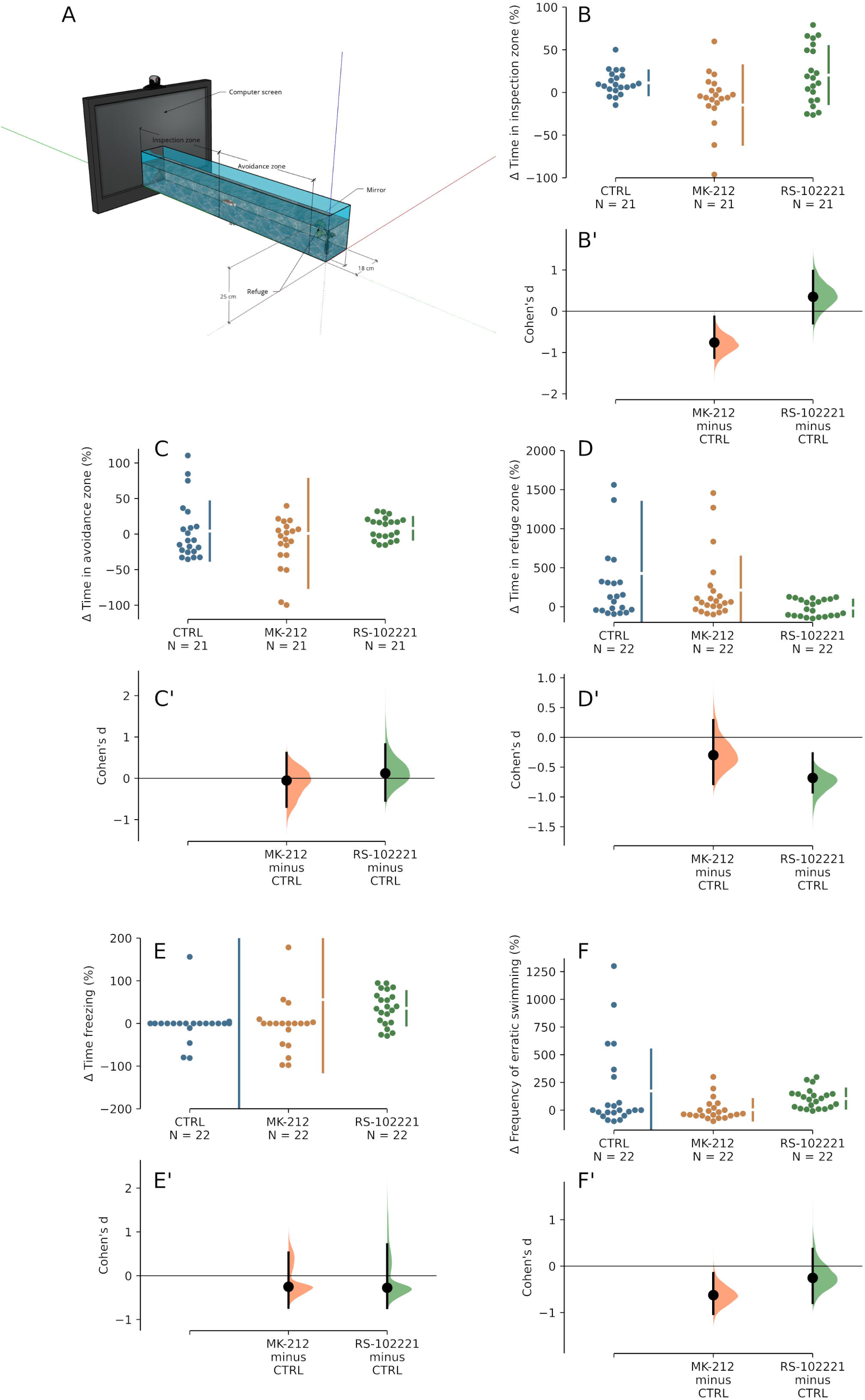
– In the conditional approach test, MK-212 decreases predator inspection and erratic swimming, while RS-102221 decreases preference for a refuge. (A) Schematics of the test apparatus. (B) Raw data for the change in time spent in the inspection zone in relation to the period before the stimulus (animated predator) was turned on. (B’) Mean differences between groups shown in (B), plotted as as bootstrap sampling distributions of Cohen’s *d* for treatments compared with controls. (C) Raw data for the change in time spent in the avoidance zone in relation to the period before the stimulus was turned on. (C’) Mean differences between groups shown in (C), plotted as as bootstrap sampling distributions of Cohen’s *d* for treatments compared with controls. (D) Raw data for the change in time spent in the refuge zone in relation to the period before the stimulus was turned on. (D’) Mean differences between groups shown in (D), plotted as as bootstrap sampling distributions of Cohen’s *d* for treatments compared with controls. (E) Raw data for the change in time spent freezing in relation to the period before the stimulus was turned on. (E’) Mean differences between groups shown in (E), plotted as as bootstrap sampling distributions of Cohen’s *d* for treatments compared with controls. (F) Raw data for change in erratic swimming in relation to the period before the stimulus was turned on. (F’) Mean differences between groups shown in (F), plotted as as bootstrap sampling distributions of Cohen’s *d* for treatments compared with controls. For all effect size plots, each mean difference (estimated Cohen’s *d*) is depicted as a dot. Each 95% confidence interval is indicated by the ends of the vertical error bars.

A mirror was positioned adjacent to the tank in a “cooperating mirror” position (that is, in parallel to the tank) (Pimentel et al., 2019, 2021), and a LCD computer screen was positioned on adjacent to the smaller side of the tank. Inside the tank, artificial plants were positioned on the first quadrant (farthest from the computer screen), providing the Refuge zone. Animals were allowed free exploration of the tank for a 5 min acclimation period, after which the video was turned on. The video displayed an animation of a sympatric predator (*Nandus nandus*), which was shown to elicit antipredator responses in zebrafish (Gerlai et al., 2009) and, when the mirror is positioned in the “cooperating mirror” position, also elicits conditional approach responses (Pimentel et al., 2021). After the video was turned on, the animal was free to explore the tank for another 10 min. Sessions were filmed from the front using a digital camera (Samsung ES68, Carl Zeiss lens). Videos were later analyzed by one of the authors (AFNP), which manually recorded the following variables, using X-Plo-Rat (https://github.com/lanec-unifesspa/x-plo-rat):

1. Change in time in the refuge zone (%): Time spent in the first quadrant of the tank after the video was turned on, as a percentage of the time spent in these quadrants before the video was turned on
2. Change in time spent in the avoidance zone (%): Total time spent in quadrants 2-6 after the video was turned on, as a percentage of the time spent in these quadrants before the video was turned on
3. Change in time spent in the inspection zone (%): Total time spent in quadrants 7-10 after the video was turned on, as a percentage of the time spent in these quadrants before the video was turned on
4. Freezing (s): Total time spent immobile, with only opercular movements being detectable, after the video was turned on
5. Erratic swimming (n): Frequency of erratic swimming movements, defined as fast and zig-zag swimming (Kalueff et al., 2013), after the video was turned on

### 2.5. Experiment 3: Role of 5-HT_2C_ receptors in mirror-induced aggressive displays in zebrafish

The mirror-induced aggression (MIA) protocol by Barbosa et al. (2019) was used. Ten animals per group were used in this experiment, based on sample size calculations on the effects of fluoxetine in zebrafish aggressive displays (Barbosa et al., 2019). After drug exposure, animals were left to rest for 30 min and then individually transferred to the test tank, a 30 cm X 22 cm X 23 cm (length X width X height) glass tank by the side of which a mirror is positioned in a 22.5° angle (Figure 4A). Animals were allowed to freely explore the tank for 3 min., after which the mirror was positioned. A video camera (Samsung ES68, Carl Zeiss lens) positioned above the tank recorded behavior for 5 min. Videos were later analyzed by one of the authors (LAM), which manually recorded the following variables, using X-Plo-Rat (https://github.com/lanec-unifesspa/x-plo-rat):

**Figure 4.**
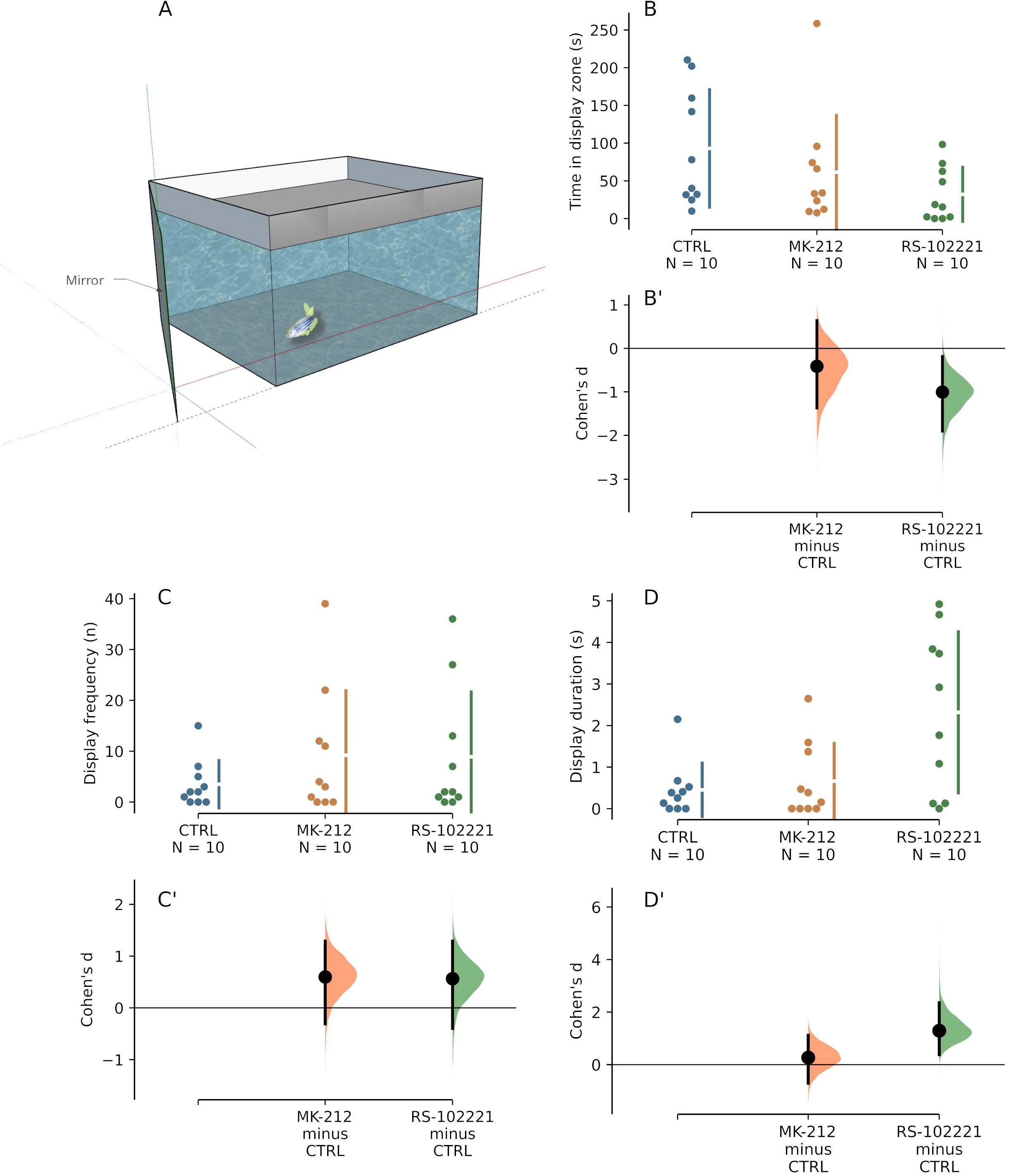
– In the mirror-induced aggressive display test, RS-102221 decreases time in display zone but increases display duration. (A) Schematics of the test apparatus. (B) Raw data for the time spent in the display zone. (B’) Mean differences between groups shown in (B), plotted as as bootstrap sampling distributions of Cohen’s *d* for treatments compared with controls. (C) Raw data for the display frequency. (C’) Mean differences between groups shown in (C), plotted as as bootstrap sampling distributions of Cohen’s *d* for treatments compared with controls. (D) Raw data for display duration. (D’) Mean differences between groups shown in (D), plotted as as bootstrap sampling distributions of Cohen’s *d* for treatments compared with controls.

1. Time spent in aggressive display (s): Total time spent in a posture that included erected dorsal, pectoral, anal, and caudal fins (Gerlai et al., 2000).
2. Frequency of aggressive displays (n)
3. Time spent near mirror (s) Total time spent in the third portion of the tank nearest to the mirror.

### 2.6. Statistical analysis

Data were analyzed based on the estimation statistics framework (Ho et al., 2019), which uses a combination of effect sizes and confidence intervals to analyze data and interpret results. Data were represented using Gardner-Altman estimation plots, with all datapoints represented as swarmplots, and effect sizes presented as a bootstrapped 95% confidence interval around Cohen’s *d* in Cumming plots (Ho et al., 2019).

## 3. Results

### 3.1. Experiment 1: Role of 5-HT_2C_ receptors in social interaction and social novelty in zebrafish

In the SI test, a medium and positive effect of MK-212 was seen on the time spent near the conspecific, while RS-102221 produced a small and negative effect on this endpoint (Figs. 1B and 1B’). No effect was seem on the time spent near the empty tank for either drug (Figs. 1C and 1C’); likewise, neither drug altered geotaxis (Figs. 1D and 1D’), erratic swimming (Figs. 1E and 1E’), or freezing (Figs. 1F and 1F’).

In the SN test, RS-102221 produced a small and negative effect on the time spent near the novel conspecific (“stranger 2”), but MK-212 had no effect (Figs. 2B and 2B’). Conversely, MK-212 produced a small to medium increase in the time spent near the known conspecific (“stranger 1”), but RS-102221 had no effect (Figs. 2C and 2C’). RS-102221 also produced a small effect on geotaxis, while MK-212 had no effect in this variable in the SN test (Figs. 2D and 2D’). Finally, no effects of either drug were observed in erratic swimming (Figs. 2E and 2E’) and freezing (Figs. 2F and 2F’).

### 3.2. Experiment 2: Role of 5-HT_2C_ receptors in conditional approach in zebrafish

A medium and negative effect of MK-212 was seen on the time spent in the inspection zone, while RS-102221 did not produce an effect on this endpoint (Figs. 3B and 3B’). No effect was found on the time spent in the avoidance zone for either drug (Figs. 3C and 3C’). RS-102221 produced a small and negative effect on time spent in the refuge zone, but no effect was found for MK-212 (Figs. 3D and 3D’). No effect was found on freezing for either drug (Figs. 3E and 3E’). Finally, a small and negative effect of MK-212 was found for erratic swimming, but no effect was found for RS-102221 (Figs. 3F and 3F’).

### 3.3. Experiment 3: Role of 5-HT2C receptors in mirror-induced aggressive display in zebrafish

A small and negative effect of RS-102221 was seen on the time in the display zone, while MK-212 did not produce an effect on this endpoint (Figs. 4B and 4B’). No effect was found on the frequency of aggressive display postures for either drug (Figs. 4C and 4C’). Finally, a small and positive effect of RS-102221 was seen on time on display (Figs. 4D and 4D’), while no effect of MK-212 was found.

## 4. Discussion

The present work evaluated the role of the 5-HT_2C_ receptor in three assays for social behavior in zebrafish: social preference (as well as social novelty), conditional approach, and mirror-induced aggression. The behavioral context for each assay is different; social preference and social novelty assesses mainly the motivation to investigate and interact with unknown conspecifics; conditional approach assesses a cooperative motivation; and the mirror-induced aggression test assesses the appetitive motivation for aggression. It was found that MK-212, a 5-HT_2C_ receptor agonist, increases preference for an unknown conspecific in the social investigation test, but also increased preference for the known conspecific in the social novelty test; the 5-HT_2C_ receptor antagonist RS-102221, on the other hand, decreased preference in the social investigation test but increased preference for the novel conspecific in the social novelty test. MK-212 also decreased predator inspection in the conditional approach test. While RS-102221 decreased time in the display zone in the mirror-induced aggressive display test, it increased display duration.

A participation of 5-HT_1A_ receptors has been proposed in zebrafish sociability; this receptor appears to be important for social preference, but not for the processing of social novelty (Barba-Escobedo and Gould, 2012). A role for 5-HT2 receptors has also been suggested in the SI test, since ritanserin, an antagonist with affinity for both the 5-HT_2A_ and 5-HT_2C_ receptor, blocked the prosocial effects of MDMA and its derivatives DOB and PMA (Ponzoni et al., 2016). In that experiment, the choice was between two images, one of a single zebrafish with the *nacre* phenotype and another of a shoal of animals of the same phenotype as the focal animal; in that situation, the focal animal is expected to prefer the shoal (Engeszer et al., 2004); MDMA, DOB, and PMA increased preference for the *nacre* image. It is not possible to infer, from Ponzoni et al.’s (2016) experiments, whether animals were reared with the presence of animals from the *nacre* phenotype, but this is an uncommon situation in zebrafish laboratories; if animals were reared in single-phenotype tanks, the novelty of the *nacre* phenotype could be an important variable to be taken into account. We found that activation of the 5-HT_2C_ receptor with MK-212 increased preference for the conspecific in the social investigation test, in consonance with Ponzoni et al.’ results; however, MK-212 increased preference for the *known* conspecific in the social novelty test, while RS-102221 increased preference for the *unknown* conspecific in this test. Thus, activation of the 5-HT_2C_ receptor appears to increase preference for conspecifics but decrease the processing of social novelty or to increase the aversiveness of novel social interactions.

Activation of the 5-HT_2C_ receptor also decreased predator inspection in the conditional approach paradigm, suggesting that this receptor is negatively involved in cooperation or altruism-like sociability. Thus, the initial hypothesis that 5-HT_2C_ receptors promote cooperative-approach behaviors has not been supported by the findings of Experiment 2. Serotonin has previously been implicated in conditional approach in guppies, with acute treatment with fluoxetine (expected to increase brain 5-HT levels) facilitating conditional approach and treatment with metergoline, a nonspecific 5-HT receptor antagonist, decreasing this behavior (Pimentel et al., 2019). The present results, however, suggest that, at least in zebrafish, cooperative-like sociability is inhibited by activation of the 5-HT_2C_ receptor. In cleaner wrasse (*Labroides dimidiatus*), the 5-HT_1A_ receptor agonist 8-OH-DPAT increased the proportion of clients inspected, while the antagonist WAY-100,635 decreased it, suggesting a role for the 5-HT_1A_ receptor in facilitating interspecific cooperation (Paula et al., 2015). Combined, these results suggest that the 5-HT_2C_ receptor has an opposite role in cooperative-like sociability in relation to the 5-HT_1A_ receptor.

Finally, while RS-102221 decreased time the animals spent in the display zone, the antagonist actually increased display duration. Serotonin has long been implicated in aggression, and pharmacological studies suggest that activation of the 5-HT_1A_ receptor inhibit aggressive display in fish (Clotfelter et al., 2007; Filby et al., 2010; Theodoridi et al., 2017). The mechanisms and receptors by which serotonin can inhibit or increase aggressive behavior in zebrafish are not yet fully understood. In mammals, the 5-HT_1A_ and 5-HT_1B_ receptors control serotonergic tone, contributing in specific brain regions, with the modulation of inhibitory postsynaptic effects on aggression (Nelson & Chiavegatto, 2001) may be regulated by the 5HT_1A_ receptor rather than the 5HT_2C_. While the role of 5-HT_2C_ receptors in aggression is not fully explored, some evidence in mice suggest that RNA editing influences not only drug efficacy, but also basal levels of aggressiveness, as VGV edited mice show increased aggression during social interactions, an effect that is rescued by treatment with MK-212 (Martin et al., 2013). de Boer et al. (2015) proposed that transient 5-HT release promotes aggressive motivation and the tonic release (basal 5-HT levels) being negatively associated with the overall level of aggressiveness, and that the 5-HT_1A_ receptor is involved mainly in that last case. The lack of effect of MK-212 in the present experiments suggest that tonic 5-HT release decreases aggression through the 5-HT_2C_ receptor as well, revealing a role of this receptor in the overall levels of aggressiveness in zebrafish.

Overall, the present results underline novel roles for 5-HT_2C_ receptors in complex social behaviors in zebrafish. Transient (phasic) activation of this receptor appears to increase social motivation, but decrease social novelty and the propensity to cooperate. Tonic serotonin, acting on 5-HT_2C_ receptors, appears to produce inhibitory effects on the processing of social novelty and aggressive display. Thus, the overall levels of serotonin, acting on this receptor, appear to negatively modulate the ability to socially appraise conspecifics. Given the general role of serotonin on emotionality and defensive behavior (Herculano and Maximino, 2014; Maximino, 2012), it is likely that basal levels of serotonin modulate the motivation to socially appraise a conspecific when non-social threats are absent, mechanisms which are mediated by the 5-HT_2C_ receptor. These mechanisms complement and balance effects that are mediated by other serotonin receptors.

## Acknowledgments

This research was made possible with the help of a productivity grant conceded to CM (CNPq, grant # 302998/2019-5).

